# Functional characterization of human Heschl’s gyrus in response to natural speech

**DOI:** 10.1101/2020.06.29.178178

**Authors:** Bahar Khalighinejad, Jose L. Herrero, Stephan Bickel, Ashesh D. Mehta, Nima Mesgarani

**Affiliations:** Mortimer B. Zuckerman Mind Brain Behavior Institute, Columbia University, New York, NY, United States; Department of Electrical Engineering, Columbia University, New York, NY, United States; Hofstra Northwell School of Medicine, Manhasset, NY, United States; The Feinstein Institutes for Medical Research, Manhasset, NY, United States

## Abstract

Heschl’s gyrus (HG) is a brain area that includes the primary auditory cortex in humans. Due to the limitations in obtaining direct neural measurements from this region during naturalistic speech listening, the functional organization and the role of HG in speech perception remains uncertain. Here, we used intracranial EEG to directly record the neural activity in HG in eight neurosurgical patients as they listened to continuous speech stories. We studied the spatial distribution of acoustic tuning and the organization of linguistic feature encoding. We found a main gradient of change from posteromedial to anterolateral parts of HG. Along this direction, we observed a decrease in frequency and temporal modulation tuning, and an increase in phonemic representation, speaker normalization, speech-sensitivity, and response latency. We did not observe a difference between the two brain hemispheres. These findings reveal a functional role for HG in processing and transforming simple to complex acoustic features and informs neurophysiological models of speech processing in the human auditory cortex.

## Introduction

Heschl’s gyrus (HG), also known as transverse temporal gyrus, is an evolutionary new part of the human auditory cortex whose functional and anatomical organization remains speculative. HG has been suggested as the location of the core area because the dense cellular structure (konicortex) and myelination of HG in postmortem tissue indicates dense thalamic input from the medial geniculate body (Brodmann, 1909; Campbell, 1905; Da Costa et al., 2011). While these architectonic studies suggest that the primary auditory cortex (PAC) is located in HG, this area is functionally heterogenous, contains multiple auditory fields (Clarke and Morosan, 2012), and has high morphological variability across individuals and brain hemispheres (Rademacher et al., 2001). The neurophysiological models of speech processing in the human auditory cortex have postulated a limited role for HG in extracting simple acoustic features (DeWitt and Rauschecker, 2012; de Heer et al., 2017; Hickok and Poeppel, 2007). Yet due to the limitations in obtaining direct neural measurements from the human auditory cortex in naturalistic listening conditions, the organization and the role of HG in speech perception remains uncertain.

Our current understanding of the functional organization of the human auditory cortex, especially in deeper cortical areas that include HG, is mostly based on fMRI neuroimaging studies. These studies have examined the organization of tonotopic, temporal modulation, and speech-sensitivity in HG. For example, modulated tones have been used to measure the temporal modulation tuning in HG, showing a medial to lateral gradient (Herdener et al., 2013; Leaver and Rauschecker, 2016; Overath et al., 2012; Schönwiesner and Zatorre, 2009). On the other hand, the lateral part of Heschl’s gyrus is found to have higher responses to speech in comparison to other sounds (Moerel et al., 2012; Norman-Haignere et al., 2015). The findings of these studies, however, have not been always consistent. For example, the proposed orientations of tonotopic map in HG include: parallel to Heschl’s gyrus (Wessinger et al., 1997), circular (Barton et al., 2012), and high-low-high gradient along the posteromedial to anterolateral axes (Dick et al., 2012; Moerel et al., 2014).

A few studies attempted to overcome the limited temporal resolution of fMRI by using direct measurement of neural activity with intracranial encephalography (iEEG). Using depth electrodes implanted in HG, these studies showed a high-low frequency tuning gradient in response to pure tones within posteromedial HG (Howard III et al., 1996; Nourski, 2017). In addition, a frequency following response (FFR) was shown to specific temporal modulations using click trains and modulated tones (Brugge et al., 2009). It is also shown that response latency increases from posteromedial to anterolateral part of HG in response to tones and isolated syllables (Nourski et al., 2014). One limitation of these studies was the use of unnatural stimuli designed to focus on a single acoustic dimension at a time (e.g. frequency or response latency). The neurons in mammalian auditory cortex, however, have multidimensional and nonlinear tuning properties (King and Nelken, 2009; Walker et al., 2011) which enable simultaneous and interactive encoding of various acoustic dimensions that occur in a naturalistic sound such as speech (Hamilton and Huth, 2018; Theunissen et al., 2000). Because the human auditory cortex responds significantly more to natural sounds than to synthetic sounds (Overath et al., 2015) and is highly specialized for speech and linguistic processing (Belin et al., 2000), several critical questions regarding the role of HG in speech processing remain unanswered. These questions include the spatial organization and multifeatured characterization of acoustic and linguistic processing in HG, the difference between tuning properties in the left and right brain hemispheres, and the role of HG in transformation of simple to complex acoustic features of speech.

To answer these questions, we used iEEG to directly measure the neural activity in a large cohort of neurosurgical patients, thus providing a comprehensive coverage of HG. We measured the neural responses as the patients listened to natural speech stories. We studied the multidimensional tuning properties of HG in response to various acoustic and linguistic attributes and measured the organization of neural responses along several acoustic dimensions to create high-resolution maps of response tuning to individual and joint acoustic and linguistic attributes. Our findings expand our knowledge of speech processing in HG and refines the current neurophysiological models of human auditory cortex.

## Methods

### Intracranial recordings

Eight adults (five females) with pharmacoresistant focal epilepsy were included in this study. All subjects underwent chronic intracranial encephalography (iEEG) monitoring at North Shore University Hospital to identify epileptogenic foci in the brain for later removal. All subjects were implanted with stereo-electroencephalographic (sEEG) depth arrays. All subjects were between 18 and 60 years old. All subjects were fluent speakers of American English and had self-reported normal hearing. In seven subjects left hemisphere and in one subject right hemisphere was dominant for language (as determined with Wada test). Electrodes showing any sign of abnormal epileptiform discharges, as identified in epileptologists’ clinical reports, were excluded from the analysis. iEEG time series were manually inspected for signal quality and were free of interictal spikes. All research protocols were approved and monitored by the institutional review board at the Feinstein Institute for Medical Research, and informed written consent to participate in research studies was obtained from each subject before implantation of electrodes.

### Data Preprocessing and Hardware

Intracranial EEG (iEEG) signals were acquired continuously at 3 kHz per channel (16-bit precision, range ± 8 mV, DC) with a data acquisition module (Tucker-Davis Technologies, Alachua, FL, USA). Either subdural or skull electrodes were used as references, as dictated by recording quality at the bedside after online visualization of the spectrogram of the signal. Speech signals were recorded simultaneously with the iEEG for subsequent offline analysis. All further processing steps were performed offline. The TDT data were resampled to 500 Hz. A 1^st^-order Butterworth high-pass filter with a cut-off frequency at 1 Hz was used to remove DC drift. Line noise at 60 Hz and its harmonics (up to 240 Hz) were removed using 2^nd^ order IIR notch filters with a bandwidth of 1 Hz. A period of silence lasting two minutes was recorded before the experiments, and the corresponding data were normalized by subtracting the mean and dividing by the standard deviation of this pre-stimulus period.

The envelope of High-gamma activity, which correlates with the neural firing in the proximity of electrodes (Buzsáki et al., 2012; Ray and Maunsell, 2011), was used as a measure of the neural response. To obtain the envelope of this broad-band signal, we first filtered the data into eight frequency bands between 70 and 150 Hz. Then, the envelope of each band was obtained by taking the absolute value of the Hilbert transform. We took the average of all eight frequency bands as the final envelope.

### Stimulus and auditory spectrogram

All stimuli were presented using a single Bose SoundLink Mini 2 speaker situated directly in front of the subject. To reduce the inevitable acoustic noise encountered in uncontrolled hospital environments, all electrical devices in patients’ room were unplugged except the recording devices and the door and windows were closed during the experiment to prevent interruption. All subjects listened to speech material containing short stories. Subjects 1 and 2 listened to stories recorded by two voice actors (one male and one female voice actor) with duration of 25 minutes and sampling rate of 16 KHz. Subjects 3 to 8 listened to stories recorded by four voice actors (two male and two female voice actors, two of the speakers were common between the two tasks) with duration of 20 minutes and sampling rate of 11025 Hz. Results were consistent when the responses to different stimuli were analyzed separately.

The time-frequency representation of speech sounds was estimated using a model of cochlear frequency analysis (Yang et al., 1992) consisting of a bank of constant 128 asymmetric filters equally spaced on a logarithmic axis. The filter bank output was subjected to a nonlinear compression, followed by a first order derivative along spectral axis (modeling inhibitory network), and finally an envelope estimation operation. This resulted in a two dimensional representation simulating the pattern of activity on the auditory nerve (Chi et al., 2005). The output of the filter bank was then resampled to 16 bands.

### Spectrotemporal receptive fields

Using the speech stimulus and high-gamma activity recorded from the implanted electrodes, we measured the spectrotemporal receptive field (STRF) of each site (Theunissen et al., 2001a). STRF is defined as a filter that predicts the neural responses from the stimulus spectrogram (Fig. 1A). STRFs were computed by normalized reverse correlation algorithm using STRFLab (Theunissen et al., 2001b). Regularization and sparseness parameters were found via cross-validation. The best frequency and response latency parameters were estimated by finding the center of the excitatory region of STRF along frequency and time dimensions. The best temporal modulation parameter was estimated from the two-dimensional wavelet decomposition of the STRF. The wavelet decomposition extracts the power of the filtered STRFs at different temporal modulations (rates) (Chi et al., 2005; Mesgarani et al., 2006). The modulation model of STRFs has four dimensions: scale, rate, time, and frequency. To estimate the best temporal modulation, we first averaged the model over three dimensions of time, frequency, and scale to calculate a rate vector. Next, we found the weighted average of the rate vector, where weights are the rate values.

**Fig1.**
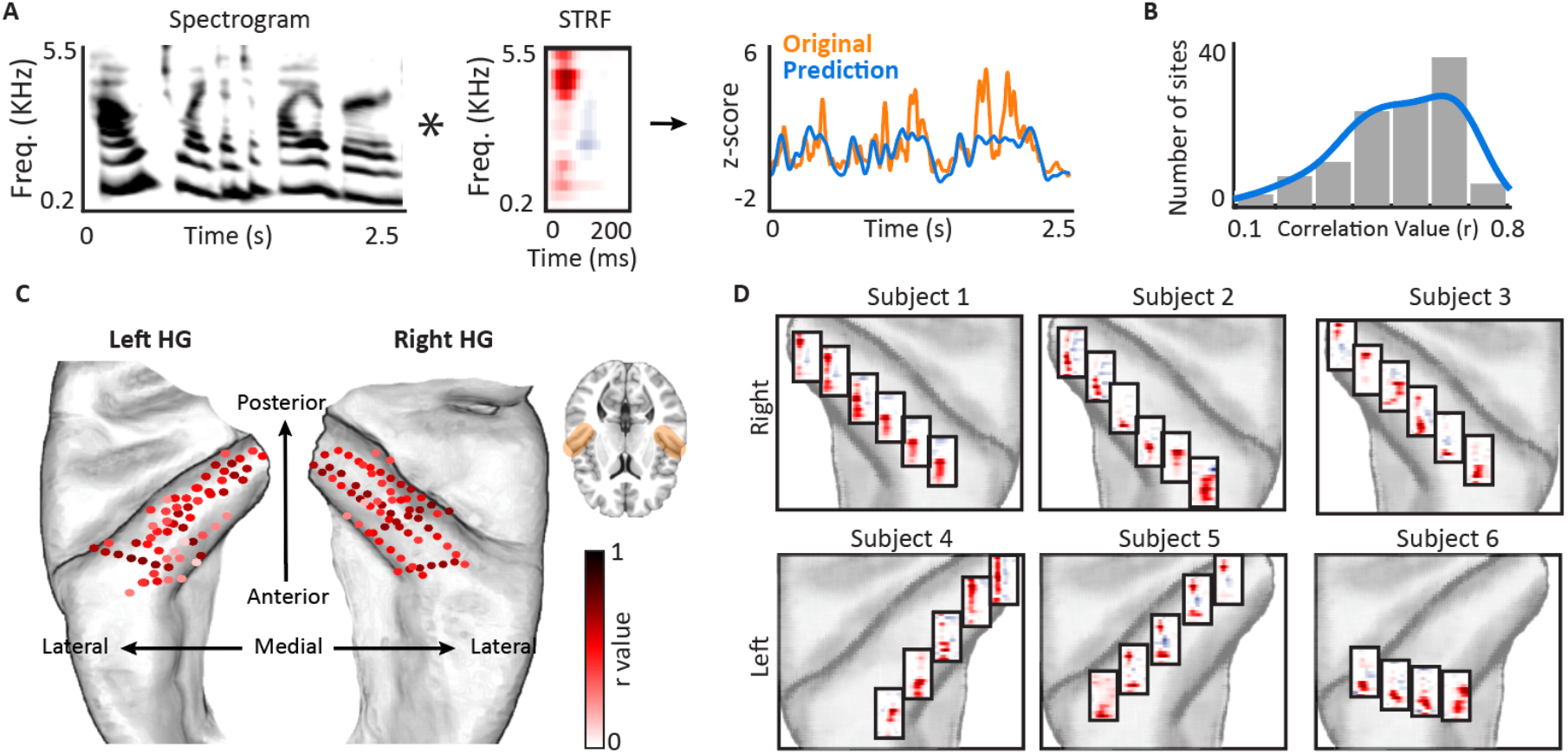
STRFs calculated from 132 sites in Heschl’s gyrus. A) Natural speech stories were played to the subjects and the spectrotemporal receptive field (STRF) was calculated for each electrode. B) The histogram of correlation values between predicted and actual responses across electrodes. C) The location of electrodes on Heschl’s gyrus and sulcus are shown on an average Freesurfer brain. The colors indicate the correlation values between predicted (20-fold cross validation) and actual response. D) STRFs of 32 example electrodes are shown for six subjects on the left (bottom row) and right (top row) Heschl’s gyrus.

### Electrodes inclusion criteria

We predicted the neural response of each site from the stimulus using the spectrotemporal receptive model (STRF, 20-fold cross-validation). The neural sites with significant prediction accuracy (Pearson correlation) were included in all subsequent analysis (t-test, false discovery rate [FDR] corrected (Benjamini and Yekutieli, 2001), p<0.01). This selection criteria resulted in 132 electrodes in Heschl’s gyrus and sulcus across all subjects (68 electrodes in right hemisphere, 64 electrodes in left hemisphere). Using these 132 electrodes, the STRFs showed an average prediction correlation of 0.44 ± 0.10 SD (Fig. 1B). The location of electrodes on an average brain is shown in Fig. 1C.

### Speech-sensitivity task

To quantify the speech-sensitivity of each neural site, six of the subjects (subjects 1, 3, 5, 6, 7, and 8) also performed a speech-nonspeech task (total of 85 electrodes). Subjects listened to 30 minutes of audio containing 69 commonly heard sounds (Supplementary figure 7). The sounds consisted of coughing, crying, screaming, different types of music, animal vocalization, laughing, syllables, sneezing, breathing, singing, shooting, drum playing, subway noises, and speech by different speakers. To determine the speech-sensitivity index, we first normalized the response of each site using the mean and variance of the neural data during the silent interval. We then averaged the normalized responses over the presentation of each sound. Finally, we performed an unpaired t-test between the averaged responses of all speech and all nonspeech sounds to obtain a t-value for each site denoting the specificity to speech over nonspeech sounds.

### Brain maps

Electrode positions were mapped to brain anatomy using registration of the postimplant computed tomography (CT) to the preimplant MRI via the postop MRI(Groppe et al., 2017). After coregistration, electrodes were identified on the postimplantation CT scan using BioImage Suite(Papademetris et al., 2006). Following coregistration, subdural grid and depth electrodes were snapped to the closest point on the reconstructed brain surface of the preimplantation MRI. We used the FreeSurfer automated cortical parcellation(Fischl et al., 2004) to identify the anatomical regions in which each electrode contact was located within approximately 3 mm resolution (the maximum parcellation error of a given electrode to a parcellated area was < 5 voxels/mm). We used Destrieux’s parcellation, which provides higher specificity in the ventral and lateral aspects of the medial lobe(Destrieux et al., 2010). Automated parcellation results for each electrode were closely inspected by the neurosurgeon using the patient’s coregistered postimplant MRI.

To calculate the topographic feature maps for each hemisphere (Supplementary figure 2), we used spatial smoothing for each tuning feature. The smoothing was done by assigning the average of the four closest neighboring electrodes to each site using knnsearch (Euclidean distance). After smoothing, a piecewise linear interpolation surface was fitted to the values of sites using Matlab fit function. In order to find the combined map for both hemispheres (Fig. 2A, Fig. 2D, Fig. 3C, Fig. 4A, Fig. 5A), the distance of each site to midsagittal plane was calculated using BioImage Suite (Papademetris et al., 2006). The absolute value of the distance to midsagittal plane was used as ML distance. For the purpose of readability, we set the minimum of ML distance to zero. The spatial smoothing was only done for visualization of the maps, but all significance tests and scatter plots were calculated using the actual raw values.

**Fig. 2.**
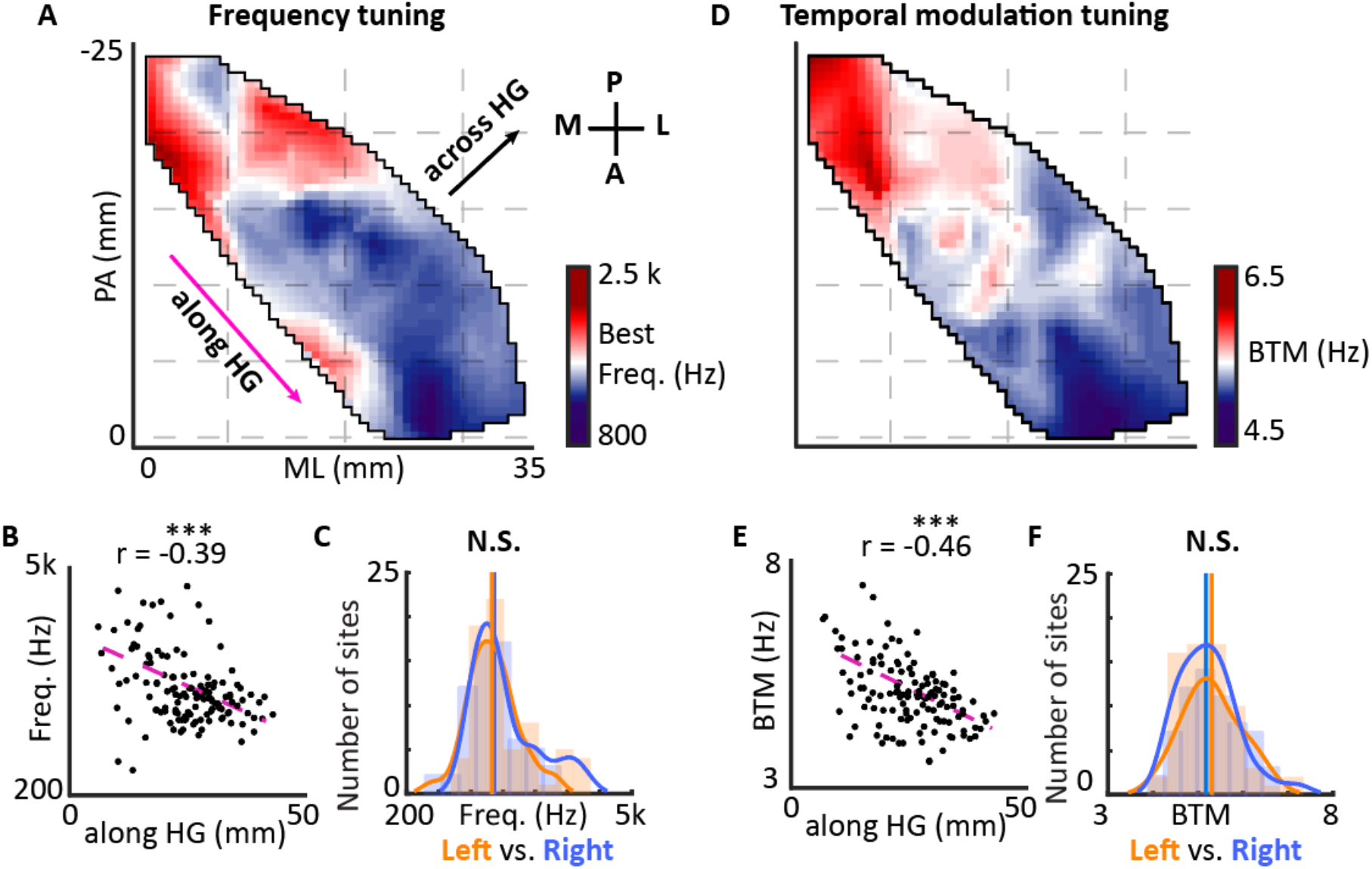
Spatial organization of frequency tuning, response latency and temporal modulation tuning. A) Spatial organization of frequency tuning is shown along two dimensions of medial to lateral (ML) and posterior to anterior (PA). B) Scatter plot of location along HG versus characteristic frequency of individual electrodes is shown (Y-axis is logarithmic). C) Histograms of characteristic frequencies estimated from neural sites in left and right Heschl’s gyrus (N.S. P > 0.1, Wilcox rank-sum test). D) Spatial organization of temporal modulation tuning. E) Scatter plot of location along HG versus temporal modulation tuning of sites. F) Histograms of temporal modulation tuning estimated from sites in left and right Heschl’s gyrus (N.S. P > 0.1, Wilcox rank-sum test).

**Fig. 3.**
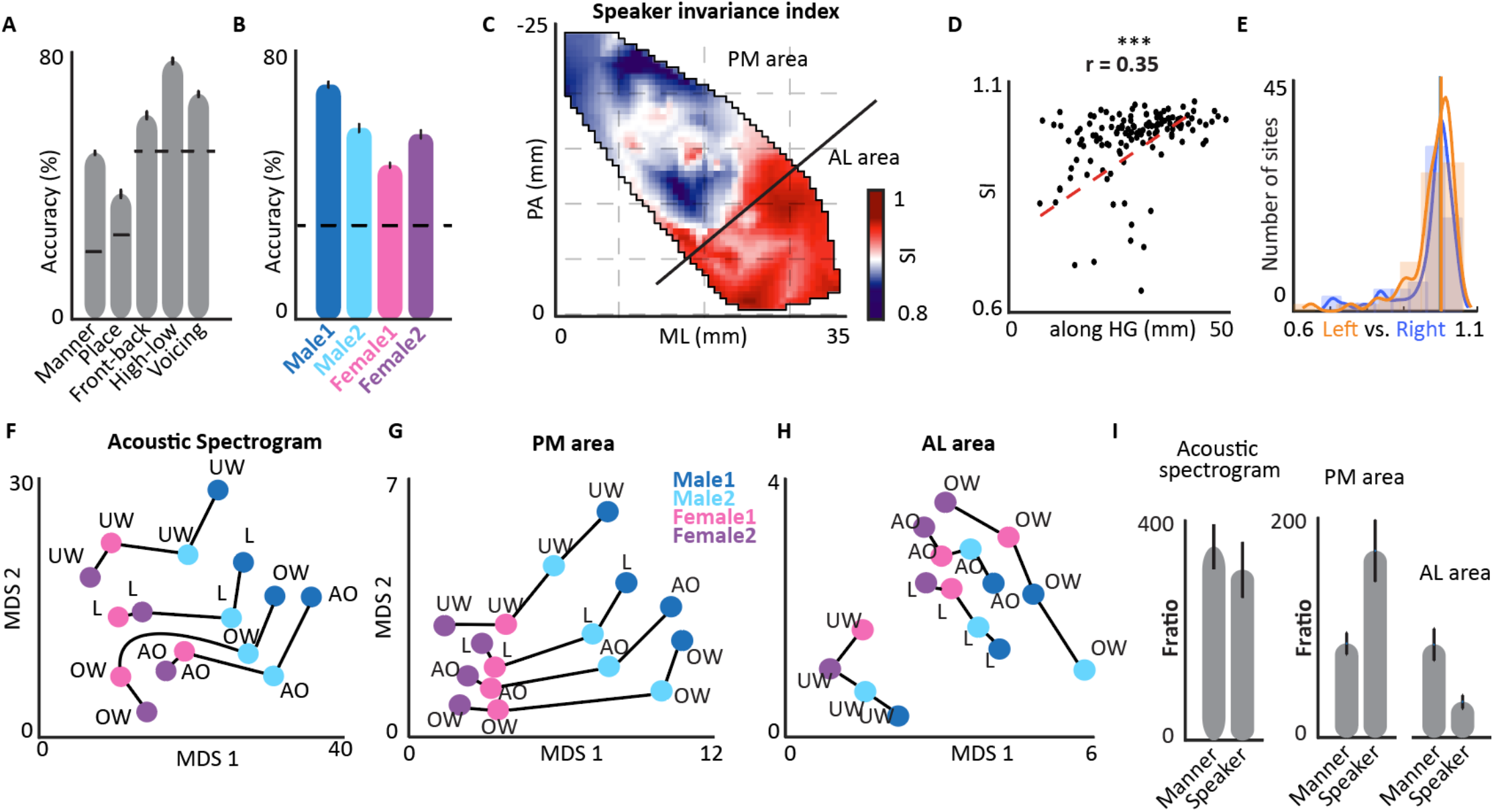
Spatial organization of speaker invariance index in Heschl’s gyrus. A) Classification accuracy of different phonetic features using population of neural sites in Heschl’s gyrus. B) Classification accuracy of different speakers using population of neural sites in Heschl’s gyrus. C) Characteristic map of speaker invariance index on two dimensions of ML and PA. D) Speaker invariance index versus location along HG for individual electrodes. E) Histograms of speaker invariance index estimated from neural sites in left and right Heschl’s gyrus (N.S. P > 0.1, Wilcox rank-sum test). F-H) MDS diagram of four phonemes spoken by four speakers derived from the acoustic spectrograms, the population neural responses in PM (posteromedial) part of HG, and in AL (anterolateral) part of HG. I) F-ratio distance between four speakers vs F-ratio distance between five manners of articulation in acoustic spectrograms, PM area, and AL area.

**Fig 4.**
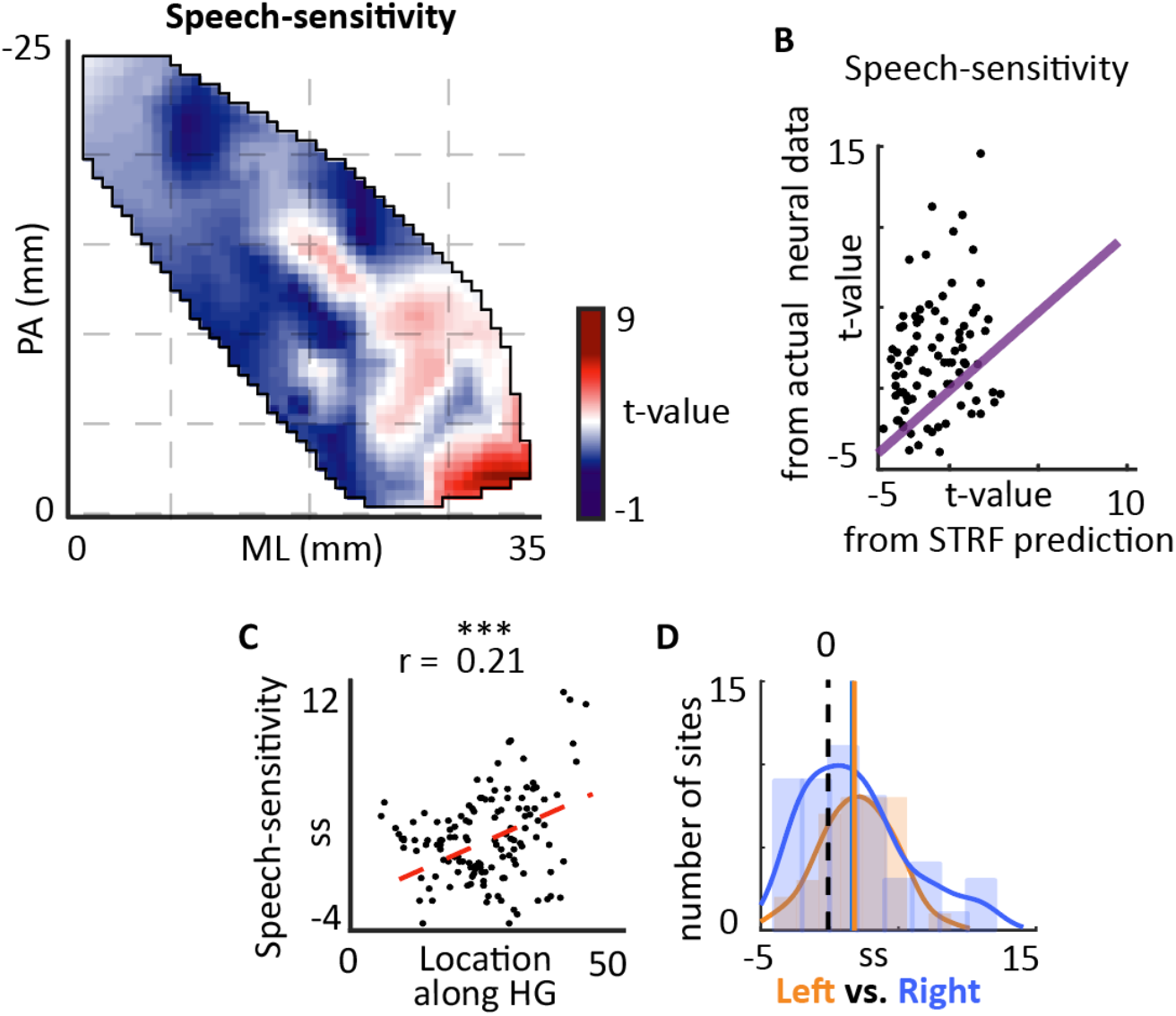
Spatial organization of speech-sensitivity. A) Spatial organization of speech-sensitivity. B) Comparison of speech-sensitivity calculated from STRF predictions versus actual neural data. C) Scatter plot of location along HG versus speech-sensitivity of individual sites. D) Histograms of speech-sensitivity estimated from neural sites in left and right Heschl’s gyrus (N.S. P > 0.1, Wilcox rank-sum test).

**Fig. 5.**
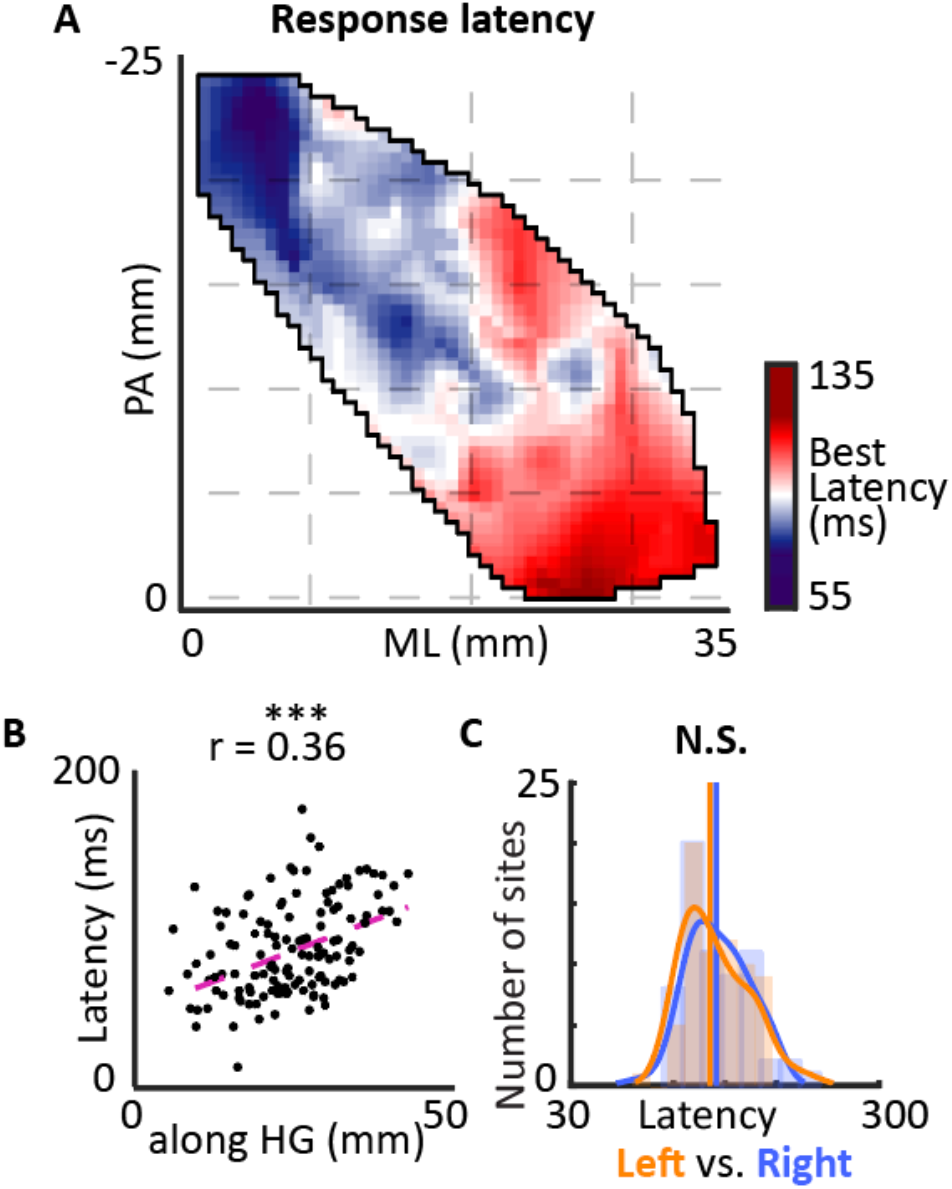
Spatial organization of response latency. A) Spatial organization of response latency. B) Scatter plot of location along HG versus response latency of individual electrodes. C) Histograms of response latencies estimated from neural sites in left and right Heschl’s gyrus (N.S. P > 0.1, Wilcox rank-sum test).

For speech-sensitivity map shown in Fig. 4A, in addition to the above steps, we also used the method of KNN imputation (N=1) to fill the values of sites for the three subjects that speech-sensitivity task was not played to. This method was only used for the purpose of visualization in Fig. 4A, and all the significance tests, correlations, and scatter plots were calculated using the actual values without KNN imputation and smoothing. The along-HG distance was calculated by projecting electrodes’ location on their first principle component (shown by purple arrow in Fig. 2A).

### Phonemes

We segmented the speech material and neural responses into time-aligned sequences of phonemes using the Penn Phonetics Lab Forced Aligner Toolkit (Yuan and Liberman, 2008). The spectrograms were aligned to the onset of phonemes with a time window of 200 ms. To minimize the preprocessing effects, we did not normalize the natural variation in phoneme length.

### MDS diagram of phonemes

To calculate the MDS diagram of phonemes for each speaker using acoustic spectrogram (shown in Fig. 3F), we first found the average acoustic spectrogram of all the instances of each phoneme spoken by each speaker. Next, the average phoneme spectrogram was windowed between 10 ms to 70 ms after the onset of the phoneme. The duration of the window was chosen according to the maximum peak of F-statistic between the categories of phonemes using all speakers (Khalighinejad et al., 2017a; Mesgarani et al., 2014). Next, we calculated the pairwise Euclidean distance between phonemes which resulted in a two-dimensional symmetric matrix reflecting a pattern of pairwise phoneme dissimilarities. To visualize this dissimilarity matrix, we used a two-dimensional multidimensional scaling algorithm (MDS) using Kruskal’s normalized criterion to minimize stress for the two MDS dimensions (Cox and Cox, 2008). The MDS diagram of phonemes based on the high-gamma responses (Fig. 3G,H) was calculated using the same method with two differences. First, each instance of a phoneme was based on the segmented high-gamma response. Second, because of the time delay between the stimuli and the response, the window was set to 90 ms to 150 ms after the onset of phoneme. This window was chosen to maximize the F-statistic of the neural responses to phoneme categories (Khalighinejad et al., 2017a; Mesgarani et al., 2014).

### Classification of phonemes and speakers

To examine the encoding of both speakers and articulation features at the population level, we trained a regularized least square (RLS) classifier (Rifkin et al., 2003) to predict the speaker or articulation feature of individual instances of PRPs (10% of data used for cross- validation). The input to the classifier was the concatenation of all the neural sites in the HG area with the window of 70 ms to 180 ms after the onset of phoneme. One classifier was trained for subjects 1 and 2, and a separate classifier was trained for subjects 3 to 8. The results of both classifiers were consistent.

### Speaker invariance index

To study the categorical encoding of phonemes in HG, we examined the similarity of the neural response to various phonemes when uttered by different speakers. Specifically, we measured the degree of speaker normalization for each neural site using a phonetic feature classifier (RLS classifier) that was trained on three of the speakers and was tested on the unseen speaker. Because the baseline decoding accuracy depends on the signal-to-noise ratio across neural sites, we divided the phonetic feature classification accuracy for the unseen speaker by the classification accuracy when the classifiers were trained within each speaker. Within speaker phoneme classification was done by training and testing of the phoneme classifier using utterances from the same speaker (10% cross-validation). This normalized phonetic feature classification accuracy on the held-out speaker was defined as the degree of phonemic categorization, and we call it speaker invariance index.

### Joint spatial organization of acoustic feature tuning

To find the correlational structure of tuning to various acoustic attributes, we used the method of principle component analysis (PCA). PCA is a dimensionality reduction technique that combines the most correlated dimensions in the data. In our analysis, each characteristic map was considered a feature and PCA was performed on *x* = (*x*_1_, *x*_2_,…,*x*_*M*_)′, where M (columns) is the number of characteristic maps, *x*_*i*_ are is a vector of N points, and N (rows) is the number of neural sites. Therefore, the PCA analysis computes the weighted sum of feature maps where weights indicate the correlation of feature maps across all neural sites. To find the dominant direction of change for each PC projection on Heschl’s gyrus, we used the canonical correlation analysis. This analysis finds the best linear combination of ML and PA distances that has the maximum correlation with the projected tuning values on the PCs. Therefore, the plotted best direction represents vector *a*, where 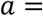 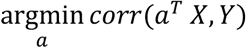, *X* is the concatenation of the ML and PA distances, and *Y* it the projected tuning values on the PCs. As a control, assigning random values to Y resulted in vectors (*a*) with random direction without preference for any specific site, confirming that the location of neural sites did not have any effect on the calculated best direction.

## Results

We used the envelope of the high-gamma frequency band (70-150 Hz) as our measure of neural response (Buzsáki et al., 2012; Ray and Maunsell, 2011). We measured five tuning attributes for each neural site: acoustic frequency, temporal modulation, speaker invariance index, speech-nonspeech response difference, and response latency. We first present the result of the tuning analysis for each of the five tuning parameters separately, and then show the properties of the joint tuning to all acoustic features. The spatial organization of each acoustic feature for all subsequent analysis is shown along the two directions of posterior to anterior (PA) and medial to lateral (ML) of HG using the location of electrodes on the average Freesurfer brain (FreeSurfer template brain, ICBM152)(Fischl et al., 2004). Across all subjects, there were 132 electrodes that were in Heschl’s gyrus and sulcus. Analyzing all subjects together provided an adequate coverage of different sections of HG along the axes of posterior to anterior (PA) and medial to lateral (ML) (Fig. 1C). We did not observe a clear spatial organization for STRF prediction accuracy (Fig. 1C). Example STRFs are shown in six of the subjects in Fig. 1D, illustrating the diversity of tuning to spectral and temporal acoustic features.

### Spatial organization of best frequency

The best frequency (BF) parameter for each cortical site is defined as the location of the excitatory peak of the STRF along the frequency dimension (Theunissen et al., 2001a). The spatial map of BF shows multiple areas with tuning to low and high frequencies, including a high-low-high frequency selective areas along HG axis, and a high to low frequency gradient on the anterolateral part of HG (Fig. 2A). Despite the mixed frequency tuning patterns in HG, we found an overall gradient of low to high frequency tuning that extends from posteromedial to anterolateral region. This gradient is shown in Fig. 2B where the distance from the posteromedial part of HG (along HG distance) is plotted against the frequency tuning for individual electrodes. The along HG distance is calculated by projecting electrodes’ location on the HG axis (shown with a purple arrow in Fig. 2A) and measuring the distance from the most medial location on HG. The high to low frequency gradient along HG is similar in both left and right HG (Supplementary figure 1,2), and there is no significant difference between the distribution of best frequency between left and right hemispheres (Wilcox rank-sum test, *P* > 0.1, Fig. 2C).

### Spatial organization of best temporal modulation

In addition to the best frequency, best temporal modulation can also be calculated from the STRF. Temporal modulation distinguishes between slowly and rapidly changing acoustic features. The human perception of sound is highly sensitive to a wide range of temporal modulations (Theunissen and Miller, 1995; Viemeister, 1979), and this acoustic attributes has shown to be an organizing factor in the human auditory cortex (Hullett et al., 2016). We defined the temporal modulation of a site as the peak of its STRF wavelet decomposition along the time axis (Woolley et al., 2005), where the transformed time axis is referred to as rate (in Hz). The difference between STRFs with different rates are shown with seven examples (slow to fast) in Supplementary figure 3. Spatial organization of temporal modulation tuning in HG is shown in Fig, 2D. The significant correlation of temporal modulation tuning with along HG distance in Fig. 2E illustrates an increase from posteromedial to anterolateral region. There is no significant difference between distribution of temporal modulations in left versus right Heschl’s gyrus (Wilcox rank-sum test, *P* > 0.1, Fig. 2F). It is also worth mentioning that the values and maps shown in figure 2 were consistent when the STRFs were estimated from two non-overlapping subset of stimulus-response pairs (test-retest), indicating the reliability of the STRF measurements across stimuli and the robustness to varying degree of neural noise (Supplementary figure 4, 5).

### Spatial organization of speaker invariance index

In previous sections we studied the organization of the best frequency and temporal modulation. However, understanding human speech is not only dependent on the acoustic attributes, but it is also dependent on the successful decoding of the linguistic units. Phonemes are the smallest contrastive units in a language and have been shown to organize the cortical responses in the human auditory cortex (Fishman et al., 2016; Khalighinejad et al., 2017a; Di Liberto et al., 2015; Mesgarani et al., 2014; Steinschneider et al., 2005). One of the major sources of acoustic variability in phones within the same phoneme category is the difference between different speakers’ voices. Normalizing speaker variability is crucial for robust decoding of the phoneme category and therefore the spoken message. At the same time, representing speaker variability is necessary for successful identification of speakers. Previous studies have shown that a categorical representation of phonemes appear in higher-level cortical areas where the encoding of phonemes becomes less sensitive to perceptually irrelevant acoustic variations (allophones) (Chang et al., 2010; Formisano et al., 2008). This emergence of phoneme categories however has not been studied in HG.

To examine the extent to which different phonemes and speaker identities are represented in HG, we used the phonetic transcription of speech utterances to obtain time-aligned neural responses to all the instances of each phoneme. we first quantified the separability of phonemes and speakers using a linear classifier trained to decode phonetic features and speaker identities (10% cross-validation was used). We restricted the analysis to five representative phonetic attributes which were the manner and place of articulation features, high-low and front-back vowel distinctions, and voiced-unvoiced attribute. We found that the population of HG responses can successfully decode all five phonetic feature significantly higher than chance (Fig. 3A). To estimate the encoding of speaker differences in HG, we classified the identity of the four speakers based on neural responses to individual phonemes. We found that the speaker differences were also decodable in the population responses with high accuracy (Fig. 3B).

To find the degree of phoneme encoding at each neural site, we defined a speaker invariance index (SI) that measures the invariance of phoneme encoding to different speakers (details in methods). We found that categorical phoneme encoding increased towards the anterolateral part of HG. Spatial organization of speaker-invariant phoneme encoding in HG is shown in Figure 3C. This figure shows two distinct encoding scheme in anterolateral and posteromedial parts of HG. The majority of sites in anterolateral HG (AL area in Fig. 3C) show a higher categorical and less speaker-dependent encoding of phonemes compared to posteromedial part of HG (PM area in Fig. 3C). This analysis shows an increase in categorical representation of phonemes towards the anterolateral part of HG. Similar to the previous maps, the phoneme encoding was also significantly correlated with the along HG distance of electrodes (Fig. 3D, Pearson correlation = 0.35, p<0.001). There was no difference between the degree of phonemic encoding and speaker normalization between left and right HG (Wilcox rank-sum test, *P* > 0.1, Fig. 3E).

To further illustrate the population encoding of phoneme and speakers in posteromedial and anterolateral HG (shown in Fig. 3C), we examined the relative distance between four representative phonemes of /UW/, /L/, /AO/, and /OW/ spoken by all four speakers. We used the first two multidimensional scaling (MDS) dimensions of phoneme responses to express their relative distances in the acoustic space and in the neural space (Fig. 3F,G,H). The population response in posteromedial HG shows a clear separation between the responses to different phonemes and speakers (Fig. 3G) similar to the representation of phonemes in acoustic space (Fig. 3F). The four speakers are shown with different colors in the MDS diagram. In contrast to posteromedial HG, the population responses in anterolateral HG (Fig. 3H) still groups the phonemes of the same category together but the separation between the phonemes of the four speakers is no longer preserved. This effect can be quantified for all phonemes and neural sites using the discriminability of the phoneme’s manner of articulations and speaker identities for acoustic spectrograms and population of sites in PM area and AL area (MDS diagram for all phonemes is shown in Supplementary figure 6). We observed that while discriminability (defined as the F-ratio (Patel et al., 1976)) of speaker identities and manners of articulation is similar in acoustic space, the discriminability of speaker identities is significantly higher than the discriminability of manners of articulation in posteromedial HG, whereas the exact opposite is true for anterolateral HG (Fig. 3I). Together these results show that the anterolateral part of HG encodes a more categorical representation of phonemes by normalizing the difference between speaker voices. In comparison, the posteromedial HG highly encodes the speaker specific differences.

**Fig. 6.**
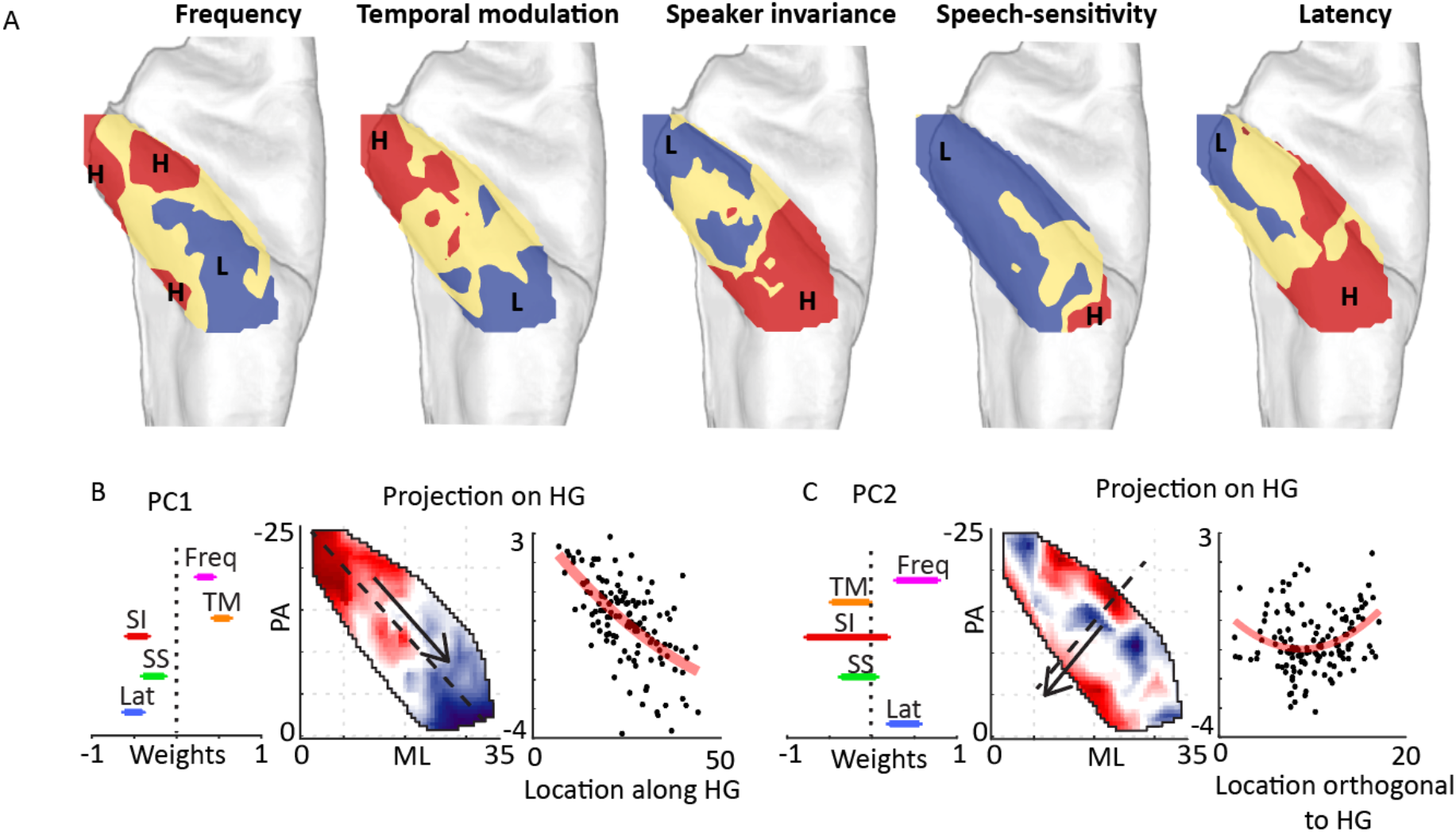
Joint spatial organization of acoustic feature tuning. Spatial organization of frequency tuning (Freq), response latency (Lat), temporal modulation (TM), speech-sensitivity (SS) and phonemic encoding (PE). B) The first principle component of joint tuning maps. The weights of the first PC are shown on the left (error bars indicate standard error calculated by bootstrapping the neural sites), projection of tuning parameters onto the first PC is shown for all sites (middle). The first PC projection versus location along HG axis is shown on the right. C) The second principle component of characteristic maps: the weights of the second PC are shown on the left (error bars indicate standard error calculated by bootstrapping the neural sites), projection of tuning parameters onto the second PC is shown for all sites in HG (middle), second PC projection versus location the orthogonal HG axis is shown on the right.

### Spatial organization of speech-sensitivity

We observed that phonemic encoding increases toward anterolateral region of HG. Since phonemes are specific to human speech, we tested whether anterolateral sites in HG also respond preferentially to speech over non-speech sounds (Chan et al., 2013; Moerel et al., 2012; Norman-Haignere et al., 2015). We define speech-sensitivity of a neural site as the t-value of a t-test between the average response of the site to speech versus nonspeech sound (Chan et al., 2013), which in our study consisted of 69 sounds (53 non-speech, 16 speech) from 14 categories (Supplementary figure 7). The speech-sensitivity values are shown on HG in Figure 4A which shows the highest values at the anterolateral side of HG. We found 39% of electrodes in HG were significantly more responsive to speech in comparison to other sounds (t-test, FDR corrected, q<0.01). To confirm that speech-sensitivity is a consequence of simple acoustic tunings and requires nonlinear transformation of the sound, we compared the actual and predicted speech-sensitivity using electrodes’ STRFs. Speech-sensitivity calculated from actual neural data is significantly higher than speech-sensitivity calculated from STRF predictions (Fig. 4B). This shows the failure of the linear STRFs to account for speech-sensitivity of sites, and confirms that this response characteristic requires nonlinear signal processing which simpler acoustic attributes such as frequency and temporal modulation tuning cannot account for.

We examined the spatial organization of speech-sensitivity using its correlation with along HG distance of electrodes. The significant correlation shown in Fig. 4C (Pearson correlation = 0.21, p<0.01) confirms a strong gradient of speech-sensitivity from the posteromedial to anterolateral part of HG. Speech-sensitivity in both hemispheres of left and right Heschl’s gyrus was significantly higher than zero, and we did not observe any difference between speech-sensitivity values in left vs. right Heschl’s gyrus (Wilcox rank-sum test, *P* > 0.1, Fig. 4D).

### Spatial organization of the response latency

The latency of the response along the auditory pathway approximately reflects the number of synapses away from the auditory periphery and hence has been used to speculate the direction of the information processing in the auditory cortex (Da Costa et al., 2011; McMurray and Jongman, 2011; Nourski et al., 2014). We defined the response latency of neural sites as the excitatory peak of the STRF along the time dimension. The observed response latencies varied from 30 to 200 ms in different parts of HG, where it was lowest in the posteromedial part of HG and gradually increased toward the anterolateral part (Fig. 5A). This gradient is shown in Fig. 5B where latency is plotted against along HG location (left vs. right is shown in Supplementary figure 1,2). There is no significant difference between the distribution of latency between the two hemispheres (Fig. 5C).

### Multivariate organization of acoustic feature tuning

Neurons in the auditory cortex have multidimensional and joint tuning to different acoustic attributes (King and Nelken, 2009; Walker et al., 2011). Our analysis so far focused on the anatomical organization of tuning to individual acoustic attributes as summarized in Figure 6A. This figure shows a correlated multidimensional organization of tuning maps to individual acoustic features. Therefore, a joint analysis of the individual tuning maps can offer further and complementary evidence for the organization of auditory fields and the main gradients of tuning change in HG.

We used an unsupervised approach to examine the organization of joint tuning and to determine the dominant anatomical directions of tuning change in HG. We performed a principal component analysis (PCA) on tuning to all five acoustic attributes across the HG sites. The PCA analysis therefore summarizes the correlation patterns among the tuning to individual acoustic attributes. We found that the first and second principal components of tuning values can account for 63% of the variance (40% and 23% respectively; third and fourth PCs are included in Supplementary figure 8). The weights of the first two PCs are shown in Figure 6B. The first PC shows that across all HG sites, a positive correlation exists between tuning to frequency and temporal modulation, which is negatively correlated with response latency, speech-sensitivity, and phoneme encoding (Fig. 6B, left). The projected tuning values on the first PC are shown in Figure 6B for all HG sites, where the dominant direction of change is calculated using the canonical correlation analysis. This analysis finds the best linear combination of ML and PA distances that has the maximum correlation with projected tuning values on the PCs. This unsupervised method shows that the direction that best describes the joint functional maps runs along the axis of HG. Because the first PC assigns significant non-zero weights to all acoustic attributes, this direction can be interpreted as the main axis along which frequency and temporal modulation tuning increase, while latency, speech-sensitivity, and speaker invariance decreases. The second PC shows the second main correlation pattern among the tuning maps and reveals a positive correlation between tuning to frequency and response latency. The projected attributes on the second PC are shown in Fig. 6C, where the direction of maximum change is orthogonal to HG direction, resulting in a secondary dominant axis of tuning gradient in HG.

## Discussion

We examined the spatial organization of multiple acoustic attributes in human HG in response to continuous speech. We found distinct spatial maps for frequency, response latency, temporal modulation, speech-sensitivity, and phonemic encoding in HG. Our results suggest that best frequency and temporal modulation tuning decreases from posteromedial to anterolateral direction in HG. On the contrary, the response latency, speech-sensitivity, and phoneme encoding increased along this HG direction. We also analyzed the properties of the joint tuning to all acoustic attributes and showed two prominent directions that explain the majority of correlated tuning changes, one along the MP-AL axis of HG and the other orthogonal to it. Compared to previous studies that used tones (Moerel et al., 2014; Nourski, 2017), ripples (Leaver and Rauschecker, 2016), or isolated phonemes (Steinschneider et al., 2011), our naturalistic speech stimuli reveals a multidimensional feature tuning in HG that organizes the responses in this auditory region.

### Organization of characteristic frequency

Tonotopy, the spatial arrangement of frequency selectivity, has been one of the fundamental organizing principles in the mammalian auditory cortex. The previous research which attempted to find the orientation of tonotopic maps in human HG is however inconclusive. Several fMRI studies that used tones and artificial stimuli showed multiple frequency-selective areas in HG. The cumulative evidence suggests that HG is located within a high-low-high frequency selective region that creates a “V” shaped pattern. Beyond this main high-low-high frequency gradient, the orientation of this tonotopy and the number of frequency selective areas has been the subject of scientific debate (Barton et al., 2012; Brewer and Barton, 2016; Da Costa et al., 2011; Dick et al., 2012; Formisano et al., 2003; Humphries et al., 2010; Moerel et al., 2012, 2014; Phillips et al., 2000; Talavage et al., 2004; Thomas et al., 2015; Upadhyay et al., 2006). Among the proposed maps of frequency tuning, our results obtained from direct recordings are most in agreement with Moerel et al. 2014 [2] which used tone pips and 7T fMRI measurement to report multiple sub-regions of low and high frequency selective areas in HG. It is worth mentioning that recording from a higher number of subjects might results in increased smoothing of frequency tuning map which may result in a V-shape low-high-low frequency selective areas similar to what has been reported by Dick et al. 2012 (Dick et al., 2012).

### Organization of temporal modulation tuning

Several previous studies have shown an encoding of temporal modulations in human auditory system (Herdener et al., 2013; Leaver and Rauschecker, 2016; Overath et al., 2012; Schönwiesner and Zatorre, 2009; Wang et al., 2011). The studies which used ripple stimuli showed that temporal rate was highest in medial HG (Herdener et al., 2013; Leaver and Rauschecker, 2016; Overath et al., 2012; Schönwiesner and Zatorre, 2009). These studies however, did not provide a precise spatial organization of temporal modulation in HG. An organized representation of temporal modulation tuning has previously been reported in the superior temporal gyrus (Hullett et al., 2016). Here, we showed that HG also has topographic representation of temporal modulation tuning which decreases from anterolateral to posteromedial HG.

### Organization of speaker invariance index

The frequency and temporal modulation tuning measures were based on a linear model of stimuli-response relationship (the STRF model) which has been commonly used to characterize the tuning of auditory cortical neurons. The higher auditory cortical regions however become progressively more nonlinear (King and Nelken, 2009). The inadequacy of linear models in such cases necessitates complementary and model-independent methods to characterize response properties. To achieve this task, we extended our linear tuning framework by examining preferential tuning to speech, and phoneme and speaker encoding. By measuring the speaker-invariance across phonetic features, we showed that speaker-invariant encoding of phonemes increases from posteromedial to anterolateral HG. This invariant encoding of phonemes suggests a processing step in creating categorical representation of phonemes in which the acoustic variability of phones imposed by different speakers is reduced. While the previous studies have shown the emergence of categorical phoneme representation in STG (Chang et al., 2010; Formisano et al., 2008; Mesgarani et al., 2014; Steinschneider et al., 2011), our results suggest that phonemic representation may be constructed in earlier areas prior to the STG. A categorical representation of phonemes involves more than just speaker normalization. Other sources of variability such as contextual and prosodic variations of phones should also be normalized to form phonemic categories. This is particularly true for more confusable allophonic variation of phonemes which may not be fully resolved in an early processing stage such as HG. Comparison of phoneme normalization in HG, PT, and STG may shed light on the progressive appearance of these linguistic units.

### Organization of speech-sensitivity

Specialization of the human auditory cortex for speech processing has been long established (Belin et al., 2000). Previous fMRI studies have shown that lateral part of HG respond more to speech compared to other sound categories and speech-like artificial stimulus (Moerel et al., 2012; Norman-Haignere et al., 2015). Our results showed that 40% of sites in HG responded preferentially to speech over non-speech sound categories, and this speech-sensitivity was highest in the anterolateral part of HG. Our observation supports the possibility that anterolateral HG might be a higher auditory field even compared to posterior STG (Nourski et al., 2014). These finding are intriguing, particularly because the cytoarchitecture studies have shown the anatomical proximity and cytoarchitectonic similarity of lateral regions of Heschl’s gyrus to the medial regions (Hackett, 2007). Further research that allows for the joint analysis of anatomical and functional properties of human HG can result in a better definition of core auditory cortex that is based on both functional and anatomical properties of the regions.

### Left and right hemisphere differences

Functional asymmetries in the human auditory cortex has long been debated in the field of neuroscience (Hickok and Poeppel, 2007). In first reports of Broca and Wernicke areas, it was shown that damage to cortical regions in left hemisphere impaired speech comprehension, but that was not the case when the damage was in the right side (Wernicke, 1874). It is also shown that lesion of the right HG disturbs the sound localization performance on both sides of space, while this was not the case for left Heschl’s gyrus (Zatorre and Penhune, 2001). Moreover, the neuroanatomy of Heschl’s gyrus shows an asymmetry where HG and PT are larger on the left side (Dorsaint-Pierre et al., 2006). In contrast to the historically established view of left lateralized speech comprehension, recent studies have argued for a bilateral involvement of the STG in speech perception and production (Bozic et al., 2010; Cogan et al., 2014). While speech can be processed bilaterally, it does not rule out the possibility of functional and computational specialization in left and right hemispheres. For example, recent studies showed differential activation of left and right hemispheres where temporal and spectral modulation processing was lateralized in left and right hemispheres accordingly (Flinker et al., 2019). It is also shown that an asymmetric processing of temporal and spectral modulation will result in an asymmetric emergence of speech and music representation in the auditory cortex (Albouy et al., 2020). It is worth noting that these studies selectively filtered out spectral and temporal modulation of speech and music, resulting in synthetic and unnatural stimuli that may activate the auditory cortex differently (Overath et al., 2015). In contrast, we did not find any difference between functional processing of left versus right HG in processing of acoustic attributes, which is consistent with proposed models of bilateral speech processing in HG (Hickok and Poeppel, 2007). Further research is needed to clarify whether the lateralization reported in previous studies (Albouy et al., 2020; Flinker et al., 2019) also occurs during naturalistic speech perception.

### Organization of response latency

The latency of the response at a neural site approximates the number of synapses that the sound has to travel from the auditory periphery before reaching that site. As such, we would expect a primary region such as auditory core area to have shorter latencies in comparison with non-primary regions such as belt and parabelt areas. In non-human primates, it has been confirmed that caudal belt and parabelt areas have shorter response latencies in comparison to rostral areas (Camalier et al., 2012; Kajikawa et al., 2005). One advantage of using direct neural measurements in our study compared to fMRI is the ability to measure the response latency with high degree of precision. We observed a wide range of response latencies in HG from 30 ms to 200 ms. While the average latency of Heschl’s gyrus is significantly lower than Superior temporal gyrus, we found that the response latency of AL-HG is comparable with, or higher than the latency of electrodes in posterior STG. This observation supports the reported findings of previous iEEG studies which used syllable and tone stimuli (Nourski et al., 2014). Similar to the frequency map, the main orientation of latency increases from MP to AL axis of HG. This result supports the notion that the primary auditory cortex is located in posteromedial part of HG. Nevertheless, the human auditory cortex is more complex than nonhuman primates with multiple core and non-core areas of auditory cortex receiving thalamic inputs from different subdivisions of the medial geniculate complex (Jones, 2003; Jones and Burton, 1976; Winer, 2010), and early activities observed in HG could potentially reflect the direct activation from auditory thalamus as opposed to the intracortical synapses (Nourski et al., 2014). As such, a conclusive separation of core vs non-core areas in humans requires simultaneous anatomical and functional analysis of HG.

### Spatial organization of joint tuning properties

While the majority of the previous research looked at the spatial organization of individual and isolated acoustic attributes, the neurons in mammalian auditory cortex have complex and multi-feature tuning properties (Bizley et al., 2013; King and Nelken, 2009; Walker et al., 2011). It is therefore crucial to examine the joint distribution of tuning properties to gain a more complete understanding of the auditory field organization, particularly because a single tuning dimension may yield ambiguous separation of auditory fields (Barton et al., 2012). Here, we adopted an unbiased and unsupervised approach to find the primary and secondary correlational structure of tuning to various acoustic attributes. Our analysis uncovered two anatomical direction. The first direction runs from medial to lateral HG and shows a gradient of change characterized by decreasing frequency and temporal modulation tuning, increasing latency, speech-sensitivity, speaker invariance, and speech-sensitivity. The second axis which was orthogonal to HG axis showed a gradient of change in frequency (low to high to low) and response latency. As such, we found both directions of medial to lateral and posterior to anterior to be important in capturing the change in multidimensional acoustic feature tuning in HG.

Relating the functional properties of neural responses in HG to the underlying anatomy remains challenging. The direction of functional change that we observed along HG is consistent with direction of anatomical gradient that are found using combined cyto- and receptor architectonic map (Morosan et al., 2001, 2005). These structural studies divided HG into three areas of Te1.1, Te1.0 and Te1.2 which extend along the HG axis. The second direction of change, orthogonal to HG is also consistent with anatomical change where Te.1 region is surrounded by Te2.1 and TI on its sides. On the other hand, while a number of structural studies have shown left dominant asymmetry in the volume of HG (Morosan et al., 2001), we did not find a difference between functional properties of left and right HGs in processing acoustic attributes. In summary, our results provide a comprehensive view of the multidimensional acoustic processing in HG and paves the way towards more complete functional characterization of auditory fields in the human auditory cortex.

## Data availability

The data that support the findings of this study are available on request from the corresponding author [N.M.].

## Code availability

The codes for performing phoneme analysis, calculating high-gamma envelope, and reconstructing the spectrogram are available at *http://naplab.ee.columbia.edu/naplib.html* (Khalighinejad et al., 2017b).

## Acknowledgments

This work was funded by a grant from the National Institutes of Health, NIDCD, DC014279, National Institute of Mental Health, R21MH114166, and the Pew Charitable Trusts, Pew Biomedical Scholars Program.

## Author contributions

B.K. and N.M. designed the experiment, evaluated results and wrote the manuscript. A.M., B.K., J.H, and N.M. collected the data. All authors commented on the manuscript.

