## Supplementary figures for "Functional characterization of human Heschl’s gyrus in response to natural speech"

**Affiliations:**

**This PDF file includes:**

Figs. S1 to S8

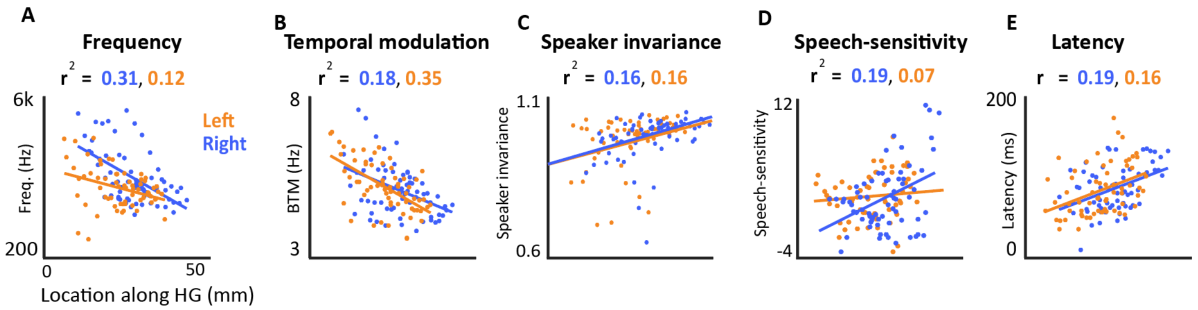


**Supplementary figure 1. Along Heschl's gyrus (HG) gradient for left vs right Heschl's gyrus**

**A to E)** Frequency, temporal modulation, speaker invariance, speech-sensitivity, and response latency for different neural sites are shown on Y-axis, the location along HG is shown on X-axis. Orange indicates the neural sites on left HG, blue indicates the neural sites on right HG.

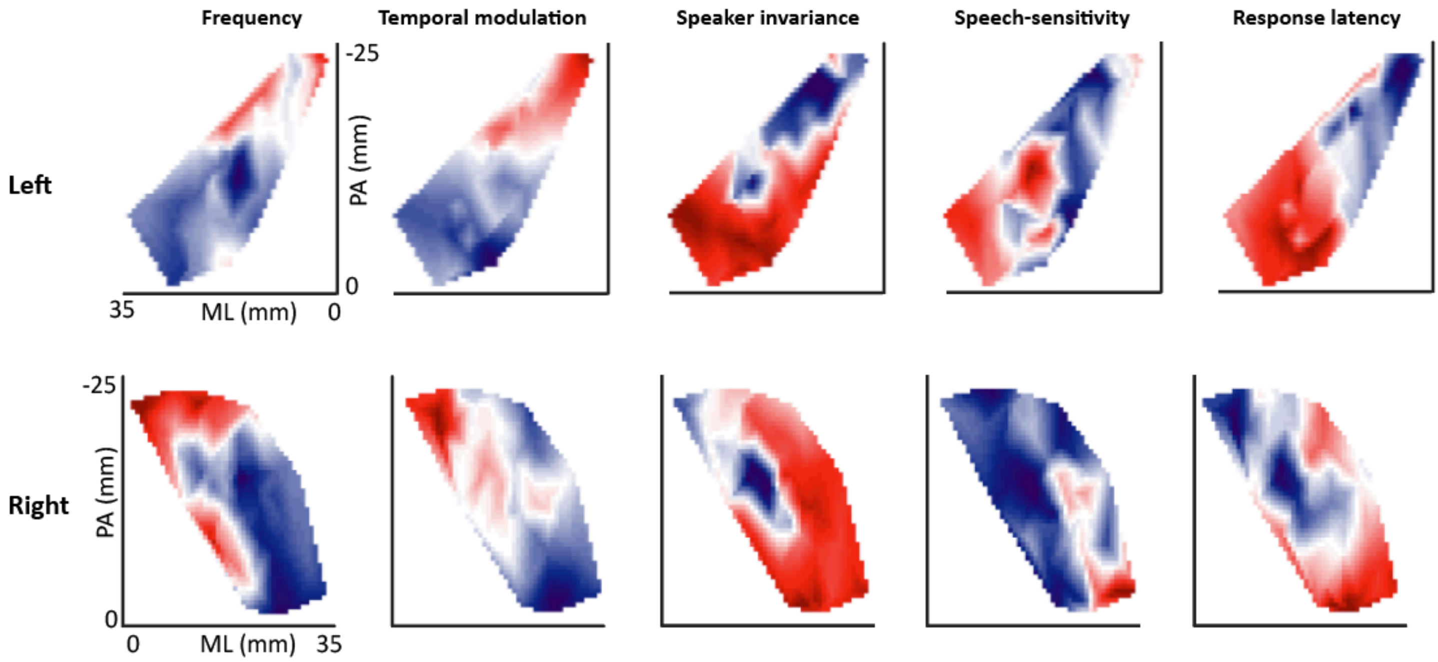


**Supplementary figure 2. Characterization maps of left and right Heschl's gyrus.**

The five characteristic maps of best frequency, best temporal modulation, speaker invariance index, speech-sensitivity and response latency are shown for left hemisphere (top row) and right hemisphere (bottom row).

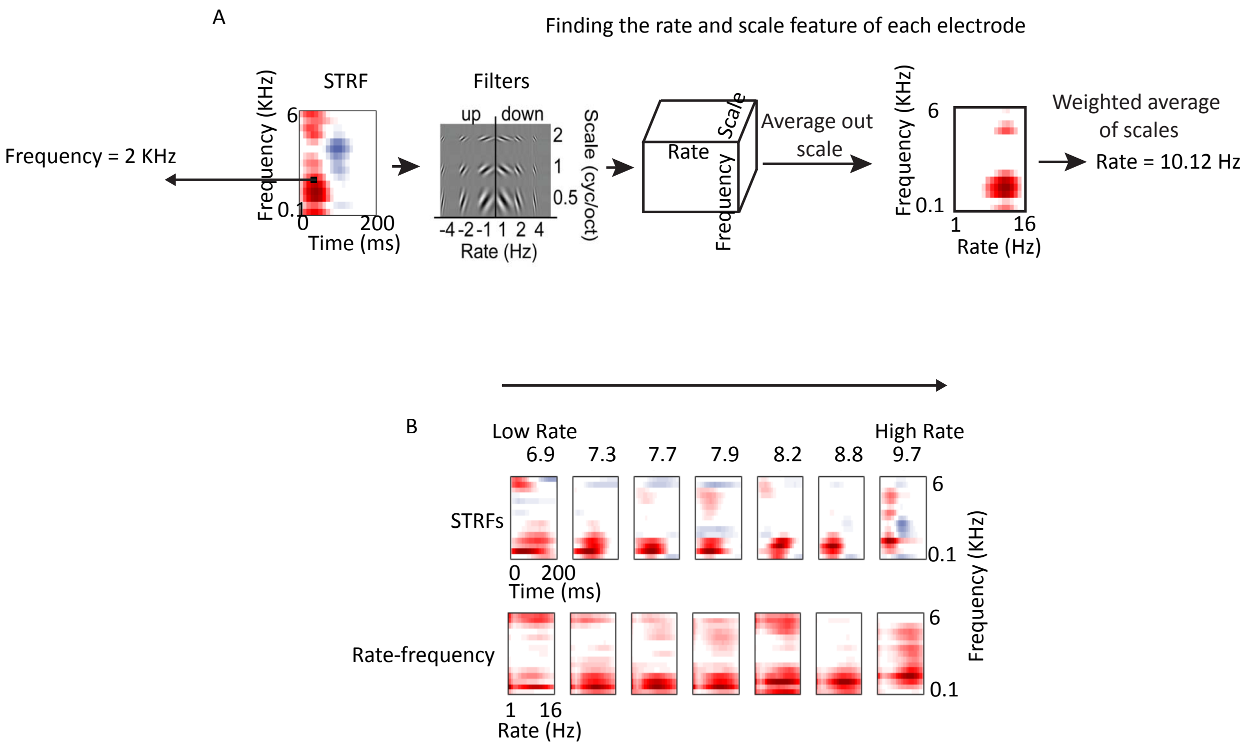


**Supplementary figure 3. Calculating rate and frequency tuning properties from STRFs.**

A) A computational model of the auditory cortex (Chi. Et. al. 2005) was used to find the temporal (rate) and spectral (scale) modulation representation of the STRFs. This transformation was done by decomposing the STRFs using 2D wavelet transforms varying along the rate and scale dimensions.

B) Example STRFs sorted based on their best rate values. STRFs with higher and lower rates are more sensitive to fast and slow acoustic changes, respectively.

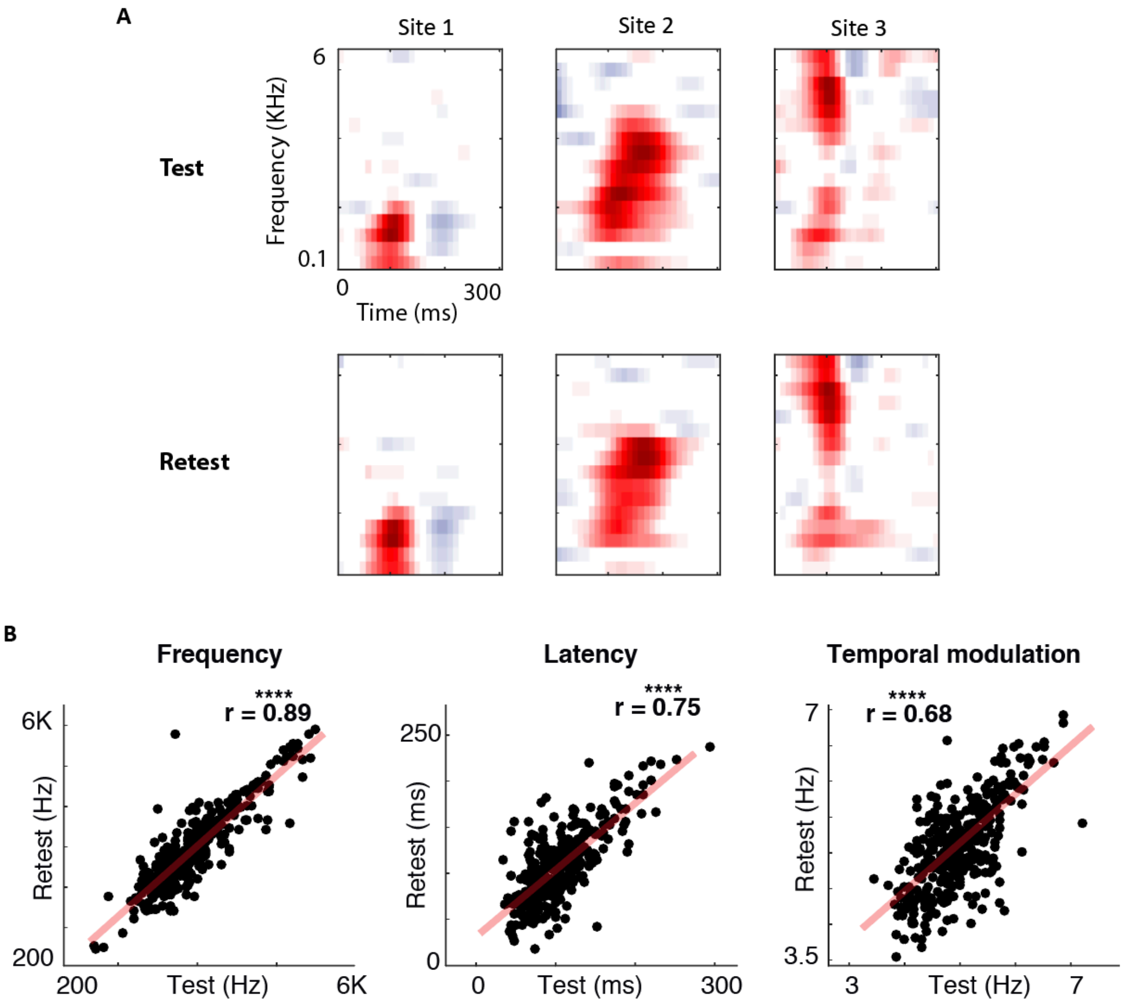


**Supplementary figure 4. Test-retest reliability of STRFs.**

Finding STRFs based on a test-retest reliability test, the stimuli and responses were segmented into two non-overlapping subsets and STRFs were calculated from each subset separately. A few example STRFs from test and retest are shown in (A) where Top row shows test and bottom row shows retest. (B) Correlation between test and retest values for best frequency, latency and temporal modulation:

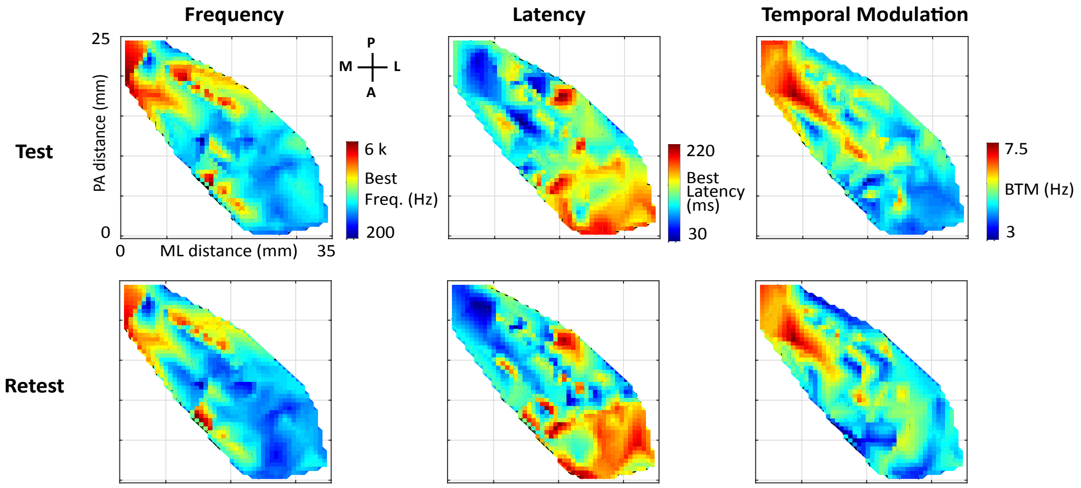


**Supplementary figure 5. Characteristic maps using test and retest stimuli.**

Characteristic maps calculated based on test (top row) and retest (bottom row) shows a high similarity between the two maps.

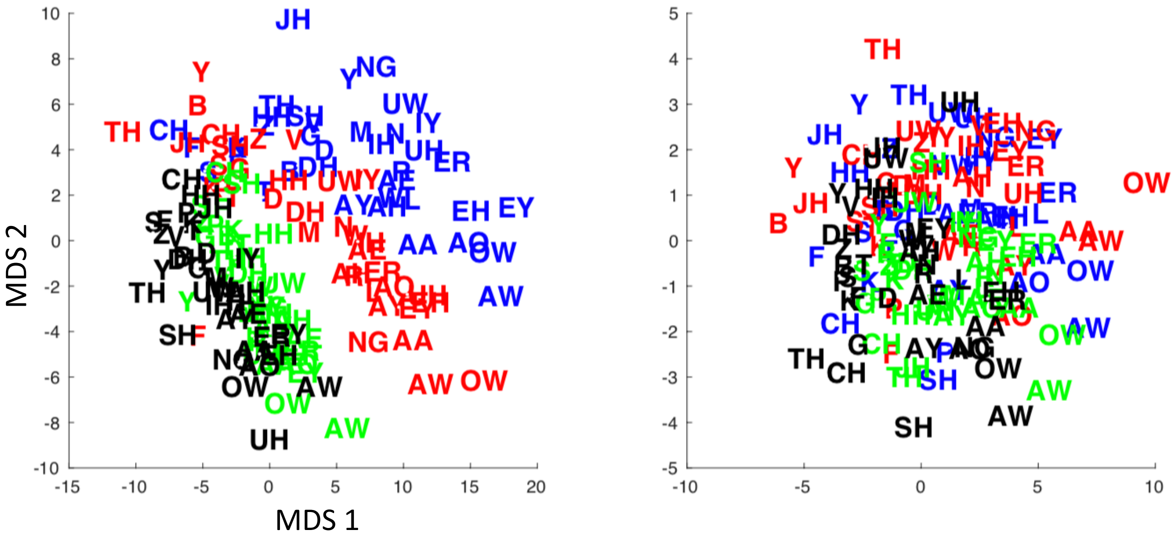


**Supplementary figure 6. MDS diagram of all phoneme for PM and AL area.**
Posteriomedial (PM) area is shown on the left, Anteriolateral (AL) area is shown on the right. The four colors indicate four different speakers where black is male 1, green is male 2, red is female 1, and blue is female 2.

A Stimulus Set: 69 commonly heard natural sounds

| | | | |
|---|---|---|---|
| 1. Classic music | 6. Speech 2 | 11. Speech 4 | 16. CV Sylablles |
| 2. Man sneezing | 7. Woman sneezing | 12. Baby crying | 17. Speech 5 |
| 3.Speech 1 | 8. Speech 3 | 13. Man laughing | 18. Drum playing |
| 4. Man breathing | 9. Woman screaming | 14. Woman sneezing | 19. Single tones |
| 5. Jazz music | 10. Pop music | 15. Gun shooting | ... |

**Supplementary figure 7. Speech-specificity task.**

**53 sounds were non-speech, 16 were speech.**

The complete list of categories of nonspeech sounds is as follows:

1. Coughing
2. Crying
3. Screaming
4. Music (Jazz, Pop, Classical)
5. Animal vocalization
6. Laughing
7. Syllables
8. Sneezing
9. Breathing
10. Singing
11. Shooting
12. Tones
13. Drum playing
14. Subway noise

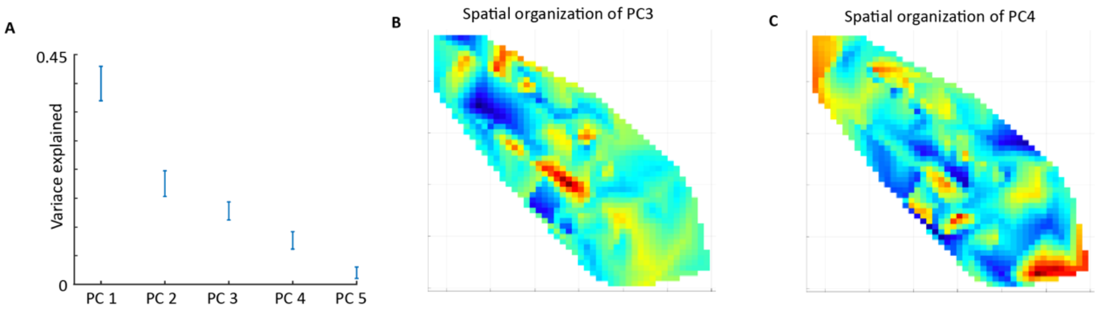


**Supplementary figure 8. Principle component analysis of joint functional characteristics.**
**A)** Variance explained by PCs 1 to 5 are shown. **B, C)** There was no spatial arrangement for PC 3 and PC 4.
